# Preventing unintentional bat captures in canopy traps used for insect surveys

**DOI:** 10.1101/2025.08.11.669606

**Authors:** Aurélien Sallé, Laurent Arthur, Guillem Parmain, David Williams, Amélie Chrétien, Elodie Le Souchu, Carl Moliard, Alain Roques, Christophe Bouget

## Abstract

There is growing interest in deploying insect traps in forest canopies for ecological research, invasive species surveillance, and biodiversity monitoring. However, forest canopies also harbor a diverse and abundant bat community, which may be incidentally trapped and killed by these devices. In this study, we investigated the impact of flight interception traps and multi-funnel traps deployed in oak canopies on incidental bat captures, and evaluated whether trap modifications could reduce this bycatch. We also examined how these modifications influenced the species richness, abundance, and mean body size of several beetle taxa, including Buprestidae, Cerambycidae, Cetoniinae, and Scolytinae. Both trap types captured bats. Flight interception traps accidentally caught seven species - *Myotis bechsteinii, Myotis daubentonii, Myotis mystacinus, Nyctalus noctula, Nyctalus leisleri, Plecotus auritus*, and *Pipistrellus pipistrellus* - with an average of 0.42 bats per trap. Multi-funnel traps primarily caught males of *P. pipistrellus*, with an average of 0.29 bats per trap. Adding a 13-mm mesh above the collection container significantly reduced bat bycatch to 0.02 and 0.05 bats per trap in flight interception traps and multi-funnel traps, respectively. This modification had no significant effect on insect species richness or mean body size, although it did reduce Cerambycidae and Scolytinae abundance in black multi-funnel and flight interception traps, respectively. This was likely due to more frequent clogging by twigs and leaves. We recommend modifying flight interception and multi-funnel traps deployed in forest canopies to prevent unintentional harm to protected bat species.

## Introduction

Insects are routinely sampled in forests for biodiversity monitoring (e.g., Kühbandner et al., 2025), invasive species surveillance (e.g., Poland and Rassati, 2019), and pest management (e.g., El Sayed et al., 2006). Such sampling provides essential information on the presence and abundance of species, helps track phenological patterns, supports predictions of pest population size, spread, and potential damage, and informs assessments of the effectiveness of silvicultural or conservation practices (Brockerhoff et al., 2023). A wide range of survey techniques - often involving traps - has been developed for detecting, monitoring, or inventorying either broad insect communities or specific target species. Sampling designs are typically tailored to optimize monitoring efficiency, and in this context, trap height can have a critical influence on trap performance (Ulyshen & Sheehan, 2017; Rassati et al., 2019). Because forest canopies offer key resources needed for the life cycle of many insect species, they can be optimal locations for trapping certain taxa (Ulyshen, 2011; Sallé et al., 2021). Placing traps in the canopy may therefore improve the detection and monitoring of both high-conservation-value insect species and major forest pests (Plewa et al., 2017; Rassati et al., 2019; Sallé et al., 2020).

Insect traps are rarely taxon-specific and frequently capture a wide range of non-target organisms. Most reported bycatch consists of invertebrates (e.g., Zumr and Stary, 1991; Rojas-Nossa et al., 2018; Hafsi et al., 2020; Sukovata et al., 2022), and in some cases, it provides an opportunity to detect rare or previously unrecorded native and exotic species (Vincent et al., 2020; Le Souchu et al., 2024a; Mas et al., 2023). However, certain trap designs can also unintentionally capture and occasionally kill vertebrates. For example, conventional pitfall traps, commonly used to sample ground-dwelling invertebrates, may also cause mortality in small mammals and amphibians (Pearce et al., 2005). The incidental capture of non-target organisms may compromise the quality and representativeness of insect samples. Decomposing bodies of vertebrates accelerate the decomposition of insect samples and may attract additional insect species (Pearce et al., 2005), while collecting a large volume of non-target organisms may also reduce trap efficiency for targeted species (Spears et al., 2021). This would in turn bias the sample and increase the time necessary to sort the samples (Pearce et al., 2005). Moreover, avoiding bycatch is important not only for ethical considerations related to the killing of beneficial and/or patrimonial species, but also for regulatory compliance, as some non-target species may be legally protected. To avoid these issues, while maintaining the effectiveness of the trap, trap designs often need to be adapted (e.g. Pearce et al., 2005; Spears et al., 2021).

Many species of bats rely on the forest canopy for roosting and foraging (Müller et al., 2013; Law et al., 2016; Erasmy et al., 2021). This is particularly true for the canopy of old-growth oak forests which harbor a large amount and diversity of microhabitats, including loose bark, cracks and cavities that are used by bats for roosting and nesting (Regnery et al., 2013). Forest insects are an important food source, and bats may play a role in the control of forest pests (Beilke and O’Keefe, 2023; Augusto et al., 2024). Bat populations in Europe are thought to have experienced significant declines in the last century, and some species have undergone significant range contractions (Browning et al., 2020). In France, eight of the 36 bat species are classified as threatened, and eight are considered near threatened (Arthur and Lemaire, 2021). In Europe, 10 of the 40 bat species are threatened (Temple and Terry, 2009), and all bat species consequently benefit from legal protection across Europe (Council Directive 92/43/EEC, 1992). Nevertheless, several studies have reported incidental captures of bats in traps intended for insect sampling (McLeod et al., 2021; Holzinger et al., 2023; Ancillotto et al., 2024). Different trap designs, including multi-funnel traps (McLeod et al., 2021; San Martin, Nyssen & Kuhn, pers. comm.), flight interception traps (Holzinger et al., 2023), adhesive traps (Ancillotto et al., 2024), MULTz traps, fan traps and yellow water traps (San Martin, Kuhn & Nyssen, pers. comm.; see Kuhn et al., 2024 for trap designs), have been reported to capture various bat species. Some authors recommended avoiding trap placement near important roosting sites of rare or sensitive species (Ancillotto et al., 2024), whilst others proposed modifications to trap designs to avoid bat bycatch (McLeod et al., 2021; Holzinger et al., 2023). However, none of these studies subsequently evaluated how such changes affected capture rates of both insects and bats.

The increasing use of insect traps for ecological research, invasive insect surveillance, and monitoring programs, along with concerns about their potential risks to bats and other non-target organisms, makes it essential to develop mitigation measures to ensure their safe use. In this study, we (i) report the incidental capture of bats by three types of insect trapping devices deployed in the canopy of oak stands in France, (ii) document the species composition and abundance of bats captured, and (iii) evaluate the effectiveness of a simple modification to these devices to minimize bat bycatch while maintaining trapping efficacy for the targeted insect groups.

## Material and methods

The study was conducted in six oak-dominated forests located in France (Table 1). The two dominant tree species in these forests were *Quercus robur* L. and *Quercus petraea* (Matt) Liebl. In each forest we selected 3 to 9 stands. The main objective of the study was to characterize the community of insects dwelling in oak canopies. For this, three kinds of traps were used: black and green multi-funnel traps (BMF and GMF respectively, Chemtica Internacional, San Jose, Costa Rica), with 12 fluon-coated funnels (Figure 1A), and flight interception traps with transparent panels (FI, Ecole d’ingénieurs de Purpan) (Figure 1B). Traps were suspended among the lower branches in the canopy (i.e. approximately 10 - 15 m above the ground). The collectors were filled with a solution of 50% (v/v) monopropylene glycol and water plus a drop of detergent. Setting GMF traps in the canopy layer is considered as one of the most efficient sampling approaches to collect *Agrilus* species (Rassati et al., 2019; Sallé et al., 2020), and can be used either to monitor native species or detect exotic ones (Santoiemma et al., 2024; Le Souchu et al., 2024b), but also to monitor and detect a wide array of canopy-dwelling insects (Sallé et al., 2020; Vincent et al., 2020; Le Souchu et al., 2024a). BMF traps were baited with volatiles (a blend of longhorn pheromones, ethanol and alpha-pinene, see Roques et al., 2023). These traps have been developed for early detection of exotic longhorn beetles (Roques et al., 2023). FI traps are routinely used to sample saproxylic beetles (Bouget et al., 2009). The traps were installed during three consecutive years, from 2019 to 2021, approximately from April to September, and were emptied every month. The number of stands, and traps per stand, varied among forests and years (Table 1).

**Table 1.**
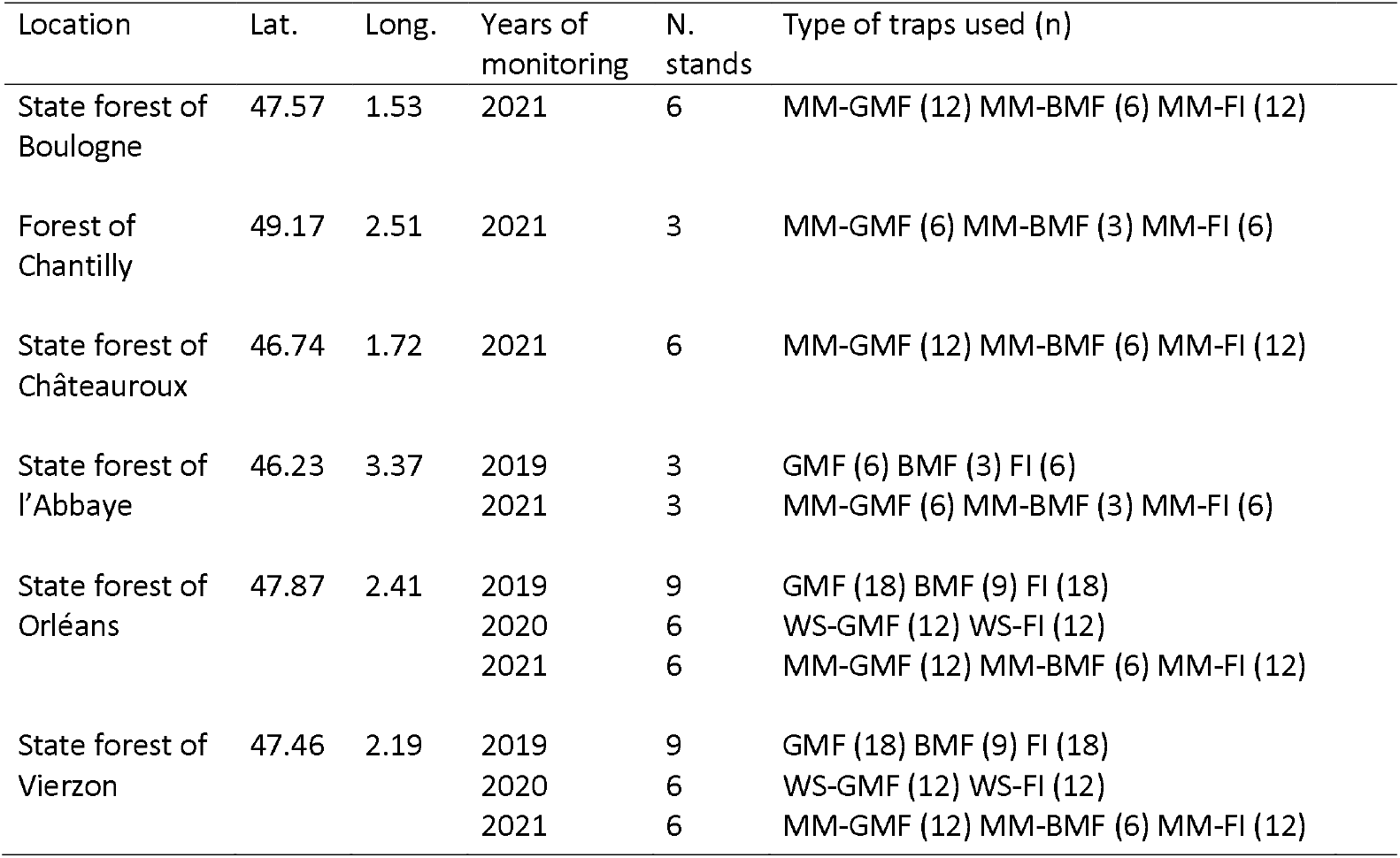
Sampling sites and years, number of stands sampled and number of traps used. Lat.: latitude, Long.: longitude, GMF: green multi-funnel traps, BMF: black multi-funnel traps, FI: flight interception traps. WS-: traps modified with wooden sticks, MM-: traps modified with a metallic mesh.

**Figure 1:**
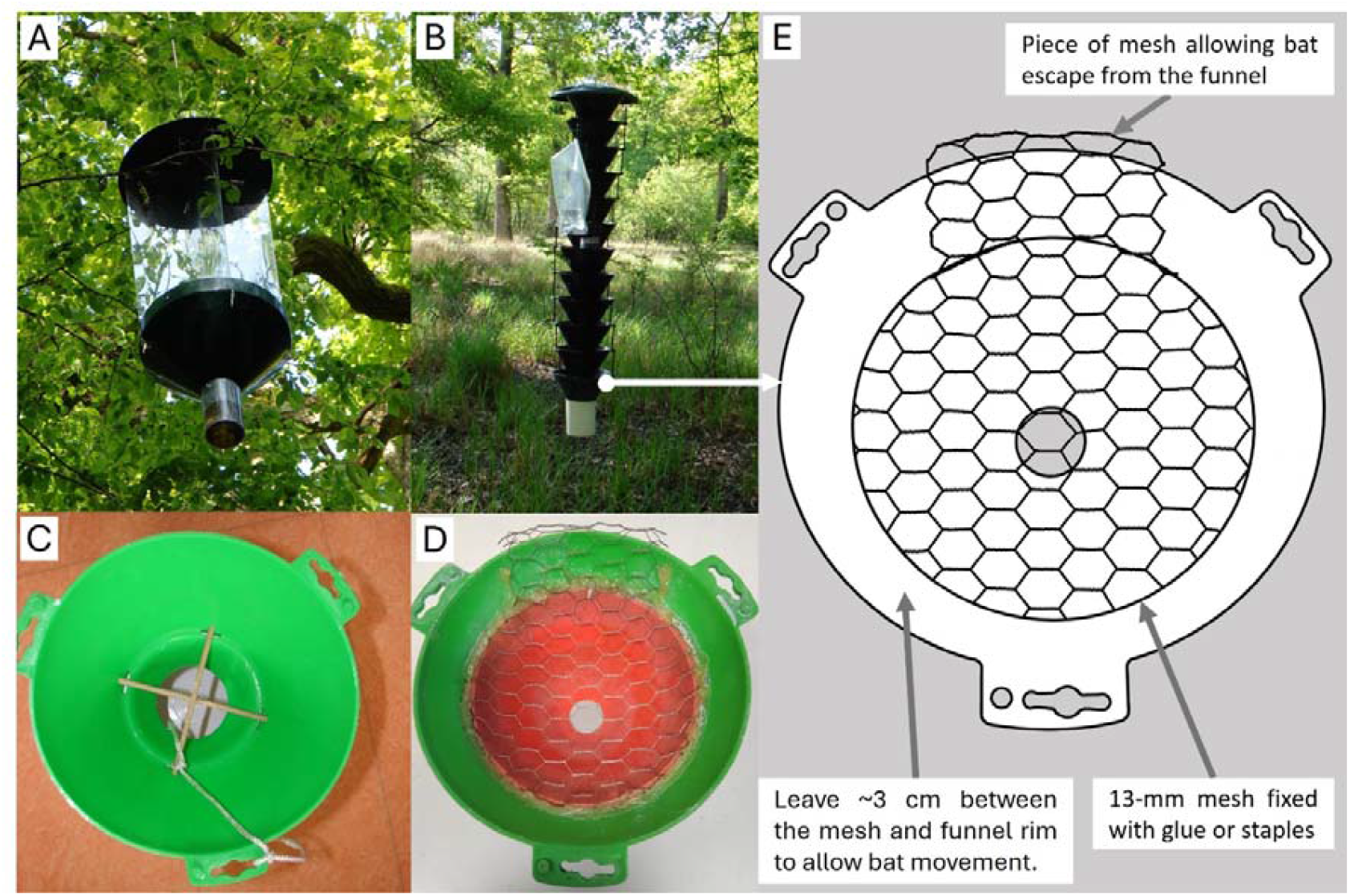
Flight interception trap (FI, A) and black multi-funnel trap (BMF, B). Modified funnels (lower funnels in both FI and BMF traps) with wooden sticks (C) and a 13-mm metallic mesh (D) placed just above the collection cup. Scheme for the modification of the last funnel (E).

In 2019, all traps were installed as supplied by the manufacturers. In 2020 and 2021, to prevent further captures of bats, as they are protected species, all traps were modified accordingly, and no unmodified (control) traps were used in parallel. In 2020, GMF and FI traps were equipped with wooden sticks (5 mm diameter), set as a cross, in the last funnel over the collection cup (Figure 1C). This modification is similar to the one proposed by Holzinger et al. (2023) to prevent bats from being captured by FI traps. BMF traps were not used in 2020. Since this modification proved ineffective (see Results), we implemented and evaluated an alternative modification in 2021. A 13-mm wire mesh was fixed onto a small funnel inserted inside the last funnel of the traps, over the collection cup of all trap types (Figure 1D and 1E), as proposed by McLeod et al. (2021). Preliminary assays conducted at the Museum of Bourges using live *Pipistrellus pipistrellus* (Schreber, 1774) from an animal shelter indicated that larger mesh sizes (2 × 2 cm and 2 × 4 cm) were easily crossed by the bats and, consequently, were too wide. The mesh was connected to the rim of the last funnel to allow bats to exit the trap (Figure 1E). We obtained legal authorization from the regional authorities (Direction Régionale de l’Environnement, de l’Aménagement et du Logement) to conduct incidental bat captures for scientific purposes under Article R411-7 of the French Environmental Code, provided that the traps were modified to reduce or prevent such captures.

### Bats and insects identification

We counted the bats caught in all types of traps during the survey. Bats caught in GMF and FI traps in 2019 and 2020 were kept for identification by the Museum of Natural History of Bourges (France). Bats collected were also sexed. Bats collected in BMF traps were counted but not identified, due to logistical constraints. Saproxylic beetles collected were identified at the species level for most families.

### Statistical analyses

All analyses were performed in R, version 4.2.1 (R Core Team 2023). To assess whether mean bat catches per trap varied among years, and after the addition of the wire mesh, we performed a Kruskal-Wallis test, followed by pairwise Wilcoxon tests, with a Holm correction for multiple comparisons.

In addition, we tested whether the change in trap design affected the abundance, species richness and average size of beetles collected. We focused these analyses on key taxonomical groups of beetles. For this, we selected several families or subfamilies of saproxylic beetles which are either significant forest pests and/or patrimonial species, exhibit various body sizes, and were well sampled by our different trap designs. Consequently we selected four groups: (i) longhorn beetles (Cerambycidae) which have average to large body sizes (13.5 ± 0.5 mm body length, calculated from the database mentioned below), include both pest and patrimonial species, and are specifically targeted by BMF traps, (ii) jewel beetles (Buprestidae) which have average body sizes (9.0 ± 0.5 mm), include pest species, and are specifically targeted by GMF traps, (iii) flower chafers (Scarabaeidae: Cetoniinae) which have large body sizes (17.4 ± 1.2 mm), include patrimonial species and are well sampled by FI traps and (iv) bark beetles (Curculionidae: Scolytinae), which have small body sizes (2.5 ± 0.1 mm), include pest species, and are also well sampled by FI traps. We assessed whether the mean abundance and species richness per trap varied before (2019) and after (2021) the addition of the wire mesh with Kruskal-Wallis tests. Because bark beetles were not the specific target of BMF traps in this study, they were not identified in these traps in 2021 and were therefore excluded from the analyses for this trap type.

Finally, to assess the effect of wire mesh on the average body size of collected beetles, we first assigned an average body size to each of the sampled taxa, within the four above mentioned taxonomic groups, using the database provided by Bouget and coll. (2019). For each group, we then calculated the average body size per trap in 2019 and 2021. For each trap type, we restricted the analyses to the most frequently sampled taxonomic groups (i.e. Buprestidae for GMF traps, Cerambycidae for BMF traps and Cetoniinae and Scolytinae for FI traps). To assess whether the average body sizes varied significantly, we performed Kruskal-Wallis tests. For all tests, the significance level was set at α = 0.05.

## Results

### Bat captures

In 2019, 33 bats were captured: 5 in the BMF traps, 11 in the GMF traps and 17 in the FI traps (Table 2). In 2020, 17 bats were caught: 8 in the GMF traps, and 9 in the FI traps. One FI trap collected 6 bats during a single survey. In 2021, 5 bats were captured: 1 in the BMF traps, 3 in one of the GMF traps and 1 in the FI traps. In 2019 and 2020, FI traps collected six species of bats, mostly *Myotis bechsteinii* (Kuhl, 1818) (37%) and *P. pipistrellus* (30%), while GMF traps collected only two species, mostly *P. pipistrellus* (95%) (Table 2). Overall, the captured specimens were predominantly male (83%). Approximately half of the captures occurred in July, with the remainder mostly distributed between May and June. While the average number of bats did not differ between 2019 and 2020 in both FI (0.42 on average) and GMF traps (0.29 on average), it significantly decreased in 2021, in all trap types (H_2, 124_ = 15.63, p < 0.001 for FI traps; H_2, 124_ = 8.06, p = 0.014 for GMF traps, H_1,50_ = 4.89, p = 0.027 for BMF traps; figure 2).

**Table 2.** Abundance of bat species captured using three types of insect traps (flight interception traps (FI), green multi-funnel traps (GMF) and black multi-funnel traps (BMF)) deployed in oak canopies. Traps operated in 2019 were used as supplied by the manufacturer (modif. no), those in 2020 were equipped with a wooden cross above the collection cup (modif. cross), and those in 2021 were fitted with a metallic mesh above the collection cup (modif. mesh). The number of traps deployed varied over years (n traps). Females (f), males (m), not determined (nd).

| Bat species |  |  |  | *Myotis bechsteinii* | *Myotis daubentonii* | *Myotis mystacinus* | *Nyctalus leisleri* | *Nyctalus noctula* | *Pipistrellus pipistrellus* | *Plecotus auritus* | not identified | Total |
| --- | --- | --- | --- | --- | --- | --- | --- | --- | --- | --- | --- | --- |
| IUCN status France/Europe |  |  |  | NT/VU | LC/LC | LC/LC | NT/LC | VU/LC | NT/LC | LC/LC |  |  |
| Trap type | Year | n traps | modif. | Number of specimens |  |  |  |  |  |  |  |  |
| FI | 2019 | 42 | no | 2 f 1 m | 2 m | 1 m | 3 m | 1 nd | 7 m |  |  | 17 |
|  | 2020 | 24 | cross | 4 f 3 m |  |  |  |  | 1 m | 1 m |  | 9 |
|  | 2021 | 60 | mesh |  |  |  |  |  |  |  | 1 nd | 1 |
| GMF | 2019 | 42 | no |  |  | 1 nd |  |  | 10 m |  |  | 11 |
|  | 2020 | 24 | cross |  |  |  |  |  | 8 m |  |  | 8 |
|  | 2021 | 60 | mesh |  |  |  |  |  |  |  | 3 nd | 3 |
| BMF | 2019 | 21 | no |  |  |  |  |  |  |  | 5 nd | 5 |
|  | 2021 | 30 | mesh |  |  |  |  |  |  |  | 1 nd | 1 |

**Figure 2:**
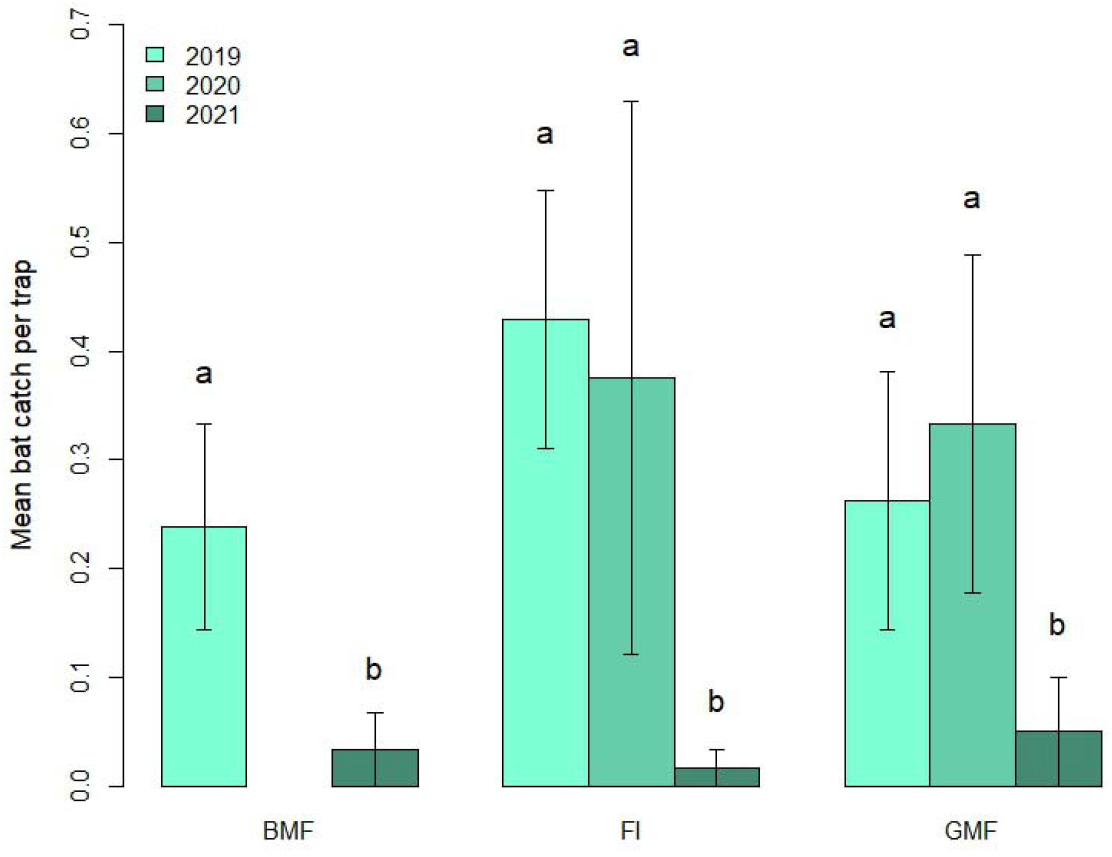
Mean number of bats (± standard error) collected in black multi-funnel traps (BMF), green multi-funnel traps (GMF), and flight interception traps (FI), in 2019 (with a standard collection cup), 2020 (with a wooden cross above the collection cup) and 2021 (with wire mesh above the collection cup). Different letters indicate statistically significant differences among groups (pairwise Wilcoxon test).

### Insect surveys

The mean abundance of beetles per trap decreased in 2021 for Cerambycidae in BMF traps and for Scolytinae in FI traps, but did not significantly vary for other taxonomical groups (*H*_1, 29_ = 6.20, p = 0.013 for Cerambycidae; *H*_1, 57_ = 4.26, p = 0.039 for Scolytinae; Figure 3). The mean =species richness per trap of the four taxonomic groups considered did not significantly vary between 2019 and 2021 (Figure 3).

**Figure 3:**
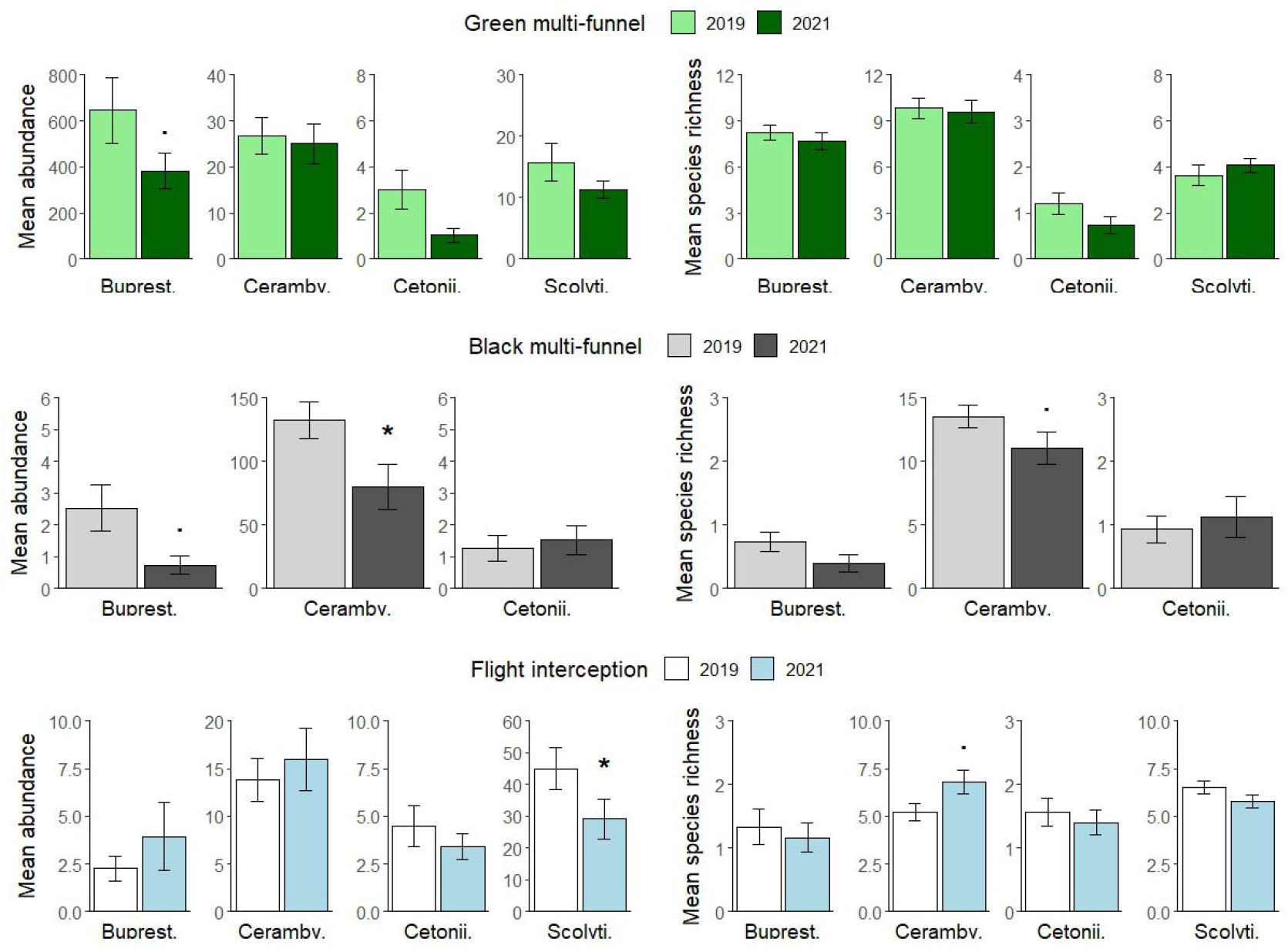
Mean abundance (± standard error) (left) and mean species richness (± standard error) (right) per trap of Buprestidae, Cerambycidae, Cetoniinae and Scolytinae collected in green multi-funnel traps, black multi-funnel traps, and flight interception traps in 2019, i.e. with a standard collection cup, and in 2021, with a wire mesh above the collection cup. Symbols: . indicates a p.value < 0.1, * indicates a p.value < 0.05. Note that abundance scales are different among families and trap types.

The average body size per trap slightly increased in 2021 for the Cerambycidae in BMF traps (*H*_1, 29_ = 4.92, p-value = 0.027; figure 4). It did not significantly vary for the other taxonomic groups considered, although it tended to decrease for the Cetoniinae (*H*_1, 48_ = 3.82, p-value = 0.051; Figure 4).

**Figure 4:**
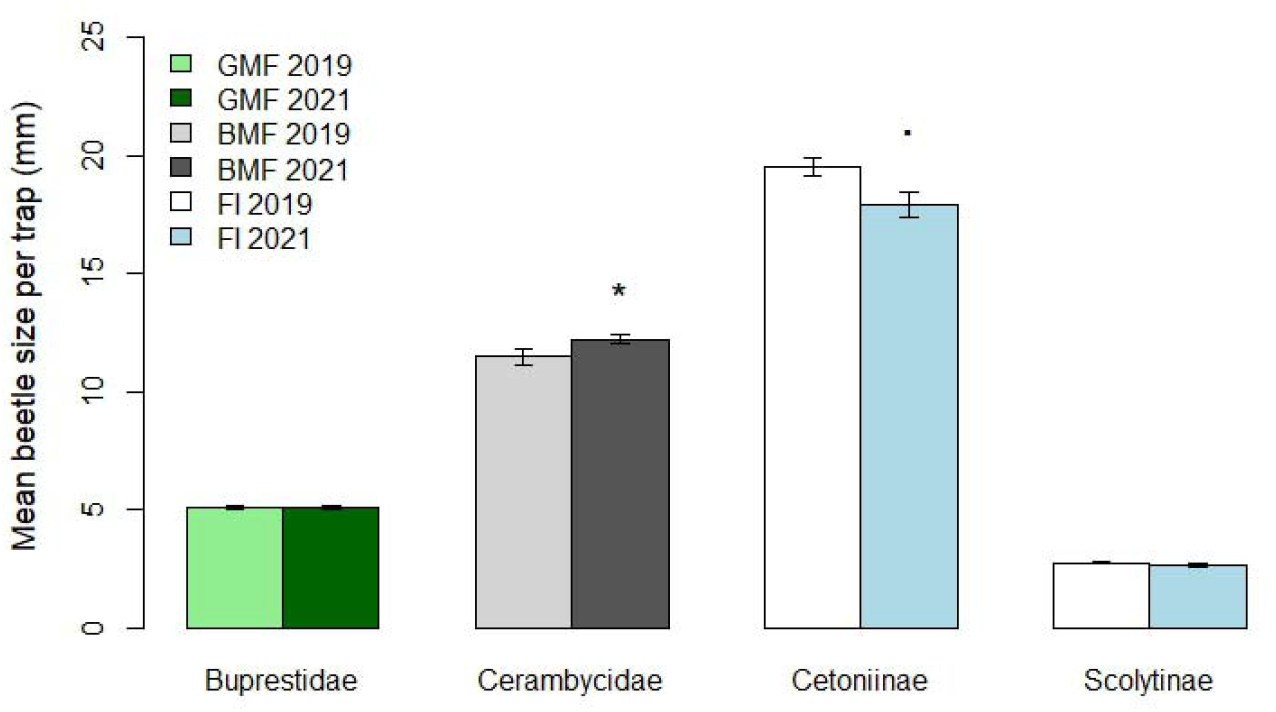
Mean body size (± standard error) per trap of species of Buprestidae, Cerambycidae, Scolytinae and Cetoniinae collected in green multi-funnel traps (green bars: Buprestidae), black multi-funnel traps (grey bars: Cerambycidae), and flight interception traps (white and blue bars: Cetoniinae and Scolytinae) in 2019, i.e. with a standard collection cup, and in 2021, with a modified collection cup. Symbols: . indicates a p.value < 0.1, * indicates a p.value < 0.05.

## Discussion

All types of trapping devices used in our study incidentally captured bats. Adding a 13-mm mesh to the funnel just above the collection cup proved to be a simple and effective modification that substantially reduces bat bycatch, without significantly affecting beetle species richness, although it may reduce overall beetle abundance.

The bat species captured in our traps are commonly found in French oak forests (Tillon et al., 2016; L. Arthur, pers. comm.), while two additional species, *Myotis alcathoe* (von Helversen & Heller, 2001) and *Barbastella barbastellus* (Schreber, 1774), are known to occur in the region but were not recorded in our survey. Comparisons with previous studies show similar patterns: FI traps in an Austrian oak forest captured *M. alcathoe, M. bechsteinii, Myotis brandtii* (Eversmann, 1845), *Nyctalus noctula* (Schreber, 1774), and *Pipistrellus pygmaeus* (Leach, 1825) (Holzinger et al., 2023), GMF traps in Belgium collected *P. pipistrellus* (San Martin, Kühn & Nyssen, pers. comm.), and in North America, BMF traps set in harvested Douglas fir stands unintentionally captured several specimens of *Myotis evotis* (H. Allen, 1864) (McLeod et al., 2021). Although most traps contained a single bat, we occasionally observed two or more individuals in the same collector, with a maximum of six female *M. bechsteinii* caught together, which explains the large standard errors in mean bat captures per trap (Fig. 2). Distress calls from trapped bats may have attracted nearby conspecifics, especially if a roost was located nearby (Russ et al., 1998). Importantly, all species we collected are protected, and the frequent capture of *M. bechsteinii* in FI traps is particularly concerning, as this species is classified as near threatened in France (Arthur & Lemaire, 2021) and vulnerable in Europe (Temple & Terry, 2009).

While FI traps captured a relatively diverse set of bat species, MF traps almost exclusively captured male *P. pipistrellus*, suggesting that these trap types differently attract or interfere with bat dispersal. Since bats use echolocation to detect prey and avoid obstacles (Jones & Holderied, 2007), it may seem surprising that they fail to avoid these large and conspicuous traps. Nevertheless, previous research has shown that bats may collide with large stationary objects (Orbach & Fenton, 2010; Greif et al., 2017; Tanalgo et al., 2025). Smooth vertical surfaces can function as ecological traps, as bats may perceive tilted smooth plates as open flyways (Greif et al., 2017). This mechanism may explain the bycatch observed in FI traps (Holzinger et al., 2023), but it does not account for captures in MF traps. One alternative explanation is that bats are attracted to insects resting on the traps or to sounds produced by trapped insects, causing them to fall into the collectors while attempting to prey on them. Another possibility, specific to MF traps, is that pipistrelles mistake these devices for potential roosting sites, resembling loose bark or bark crevices that they commonly use for summer roosts (Bat Tree Habitat Key, 2018; Holzinger et al., 2023). In our study, most pipistrelle captures occurred during the maternity and juvenile-rearing period (May–July). During this time, colonies are predominantly composed of females, which typically roost in anthropogenic structures within towns and villages (Mathews et al., 2023). Solitary males, by contrast, tend to forage in more distinct areas, including forests (L. Arthur, pers. obs.), which likely explains why males overwhelmingly dominated our captures as well as those reported in Austria (Holzinger et al., 2023) and Belgium (San Martin, Kühn & Nyssen, pers. comm.).

Installing a 13-mm metallic mesh significantly reduced the number of bat captures. Because standard and modified traps were not deployed simultaneously, due to concerns about further bat mortality, we cannot entirely rule out that the reduction observed in 2021 was influenced by interannual variation in bat populations. However, this explanation seems unlikely, as it would require a concurrent decline across multiple species. Moreover, the fact that modified traps have since been used in forest canopies in France, the United Kingdom and Belgium with very few recorded bat captures further supports the effectiveness of the mesh. The mesh did not eliminate bat captures entirely, as a few individuals were still collected in 2021. Close inspection of modified traps suggests that the gap between the mesh and upper funnel may have been too narrow, preventing bats from escaping and forcing them downward through the mesh. This highlights the need for careful installation of the mesh. Using a finer mesh might further reduce bat bycatch, though it could also impede insect capture. Additionally, a square mesh may be preferable to a hexagonal one, as bats’ forelimbs might become trapped in the latter (A. Chrétien, pers. obs.). In contrast, the addition of wooden sticks did not prevent bats from entering the collectors, indicating that the modification proposed by Holzinger et al. (2023) is unlikely to effectively reduce bat bycatch. The gaps between the sticks were probably too large to stop bats from falling into the collection cup, and bats likely require a ladder-like structure to escape from the final funnel. While adding a piece of mesh appears to be an effective measure, the rope installed with the wooden cross was insufficient, highlighting the need for careful trap design to minimize bat captures.

The trapping devices used in this study are commonly employed for insect detection and monitoring, and the use of MF traps is expected to increase in Europe in response to biological invasions. MF traps were originally developed for bark beetle sampling (Lindgren, 1983) and are now widely used globally for forest insect surveillance programs. For instance, BMF traps are widely used for biosurveillance of wood-boring insects at points of entry such as ports and airports (Roques et al., 2023). FI traps are also routinely used for forest insect monitoring (Bouget et al., 2009). However, these traps are typically deployed at ground level, where interactions with bats may be less frequent. This likely explains the limited number of previous reports of bat captures in these trap types. Since the introduction of the emerald ash borer (*Agrilus planipennis* Fairmaire, 1888) in North America, GMF traps have been extensively used for its detection and monitoring (Crook et al., 2014; Santoiemma et al., 2024). These traps are ideally placed in the canopy (Crook et al., 2014), as newly emerged adults feed and mate in the foliage. *Pipistrellus pipistrellus* is absent from North America but widespread in the Western Palearctic (Mathews et al., 2023), which may explain why, despite extensive MF trap use in North America, large-scale bat bycatch has not been reported. The emerald ash borer has also been introduced into Russia and is spreading westward across Europe, posing a major threat to native ash populations (Orlova-Bienkowskaja, 2014; Orlova-Bienkowskaja et al., 2020). In Europe, deploying GMF traps in ash canopies is recommended for early detection (European Food Safety Authority et al., 2020; Evans et al., 2020; Santoiemma et al., 2024), enabling timely management interventions. However, if not properly adapted, our results indicate that widespread deployment of GMF traps across Europe could inadvertently cause significant bat mortality, including among species classified as vulnerable in France (e.g., *N. noctula*) or Europe (e.g., *M. bechsteinii*).

Adding a mesh could potentially reduce both the quantity and diversity of insects captured, particularly larger individuals capable of escaping before falling into the collector (Pearce et al., 2005). In the context of biosurveillance, maximizing trap capture numbers is critical because exotic species ideally need to be detected when the population is at a very low abundance. While adding a mesh reduced the numbers of individuals of some species captured in our study, we did not observe any significant reduction in mean species richness, hence this implies that trap efficacy at detecting quarantine or other target exotic species is unlikely to be detrimentally affected by the addition of the mesh. Similarly, there was no significant decrease in mean beetle body size, even among larger taxa such as Cerambycidae and Cetoniinae. We did, however, note significant reductions in abundance for some groups (Cerambycidae and Scolytinae) in specific trap types in 2021 when the mesh was installed, with a similar trend observed for Buprestidae. As with bat captures, because standard and modified traps were not used simultaneously, we cannot conclusively attribute these decreases to the mesh, particularly given the well-documented interannual variability in forest insect populations (e.g., Garnas et al., 2023; Le Souchu et al., 2024b). The reduced abundance may also result from mesh clogging by foliage or other debris, which could block insects from reaching the collector; more frequent maintenance could help mitigate this issue.

Our results confirm that FI and MF traps incidentally capture bats, including species of high conservation concern in France and Europe, such as *M. bechsteinii, N. leisleri, N. noctula*, and *P. pipistrellus*. We show that a simple modification, i.e. adding a 13-mm mesh above the collection cup, can substantially reduce bat bycatch while having minimal impact on beetle species richness, though it may lower overall beetle abundance. Further dedicated studies should simultaneously evaluate modified and unmodified traps, with the use of dry collection cups and daily inspections to reduce the risk of harm to bats in unmodified traps. We recommend that all traps deployed in forest canopies, at least in Europe, be systematically modified to prevent unintentional harm to bats, and we urge trap manufacturers to integrate this consideration into future designs.

## Acknowledgements

This work was supported by the Région Centre-Val de Loire Project no. 2018-00124136 (CANOPEE), coordinated by A. Sallé. We thank O. Denux (INRAE) and X. Pineau (University of Orléans) for their technical assistance. We also thank G. San Martin (CRAW), A. Kühn (CRAW), and P. Nyssen (Ecofirst sc) for providing data on bat captures from their various entomological surveys in Belgium. Finally, we are grateful to the reviewers for their constructive comments. The DREAL provided legal authorizations for the incidental capture of bats, according to article R411-7 of the French Environmental Code (# 41-2021-04-26-00004, 36-2021-04-07-00008, DDT-2021/001).

